# Unraveling the Structure of Meclizine Dihydrochloride with MicroED

**DOI:** 10.1101/2023.09.05.556418

**Authors:** Jieye Lin, Johan Unge, Tamir Gonen

**Affiliations:** Department of Biological Chemistry, University of California, Los Angeles, 615 Charles E. Young Drive South, Los Angeles, California 90095, United States; Department of Physiology, University of California, Los Angeles, 615 Charles E. Young Drive South, Los Angeles, California 90095, United States; Howard Hughes Medical Institute, University of California, Los Angeles, Los Angeles, California 90095, United States

## Abstract

Meclizine (Antivert, Bonine) is a first-generation H1 antihistamine used in the treatment of motion sickness and vertigo. Despite its wide medical use for over 70 years, its crystal structure and the details of protein-drug interactions remained unknown. In this study, we used microcrystal electron diffraction (MicroED) to determine the three-dimensional (3D) crystal structure of meclizine dihydrochloride directly from a seemingly amorphous powder. Two racemic enantiomers (R/S) were found in the unit cell, which packed as repetitive double layers in the crystal lattice. The packing was made of multiple strong N-H···Cl^-^ hydrogen bonding interactions and weak interactions like C-H···Cl^-^ and pi-stacking. Molecular docking revealed the binding mechanism of meclizine to the histamine H1 receptor. A comparison of the docking complexes between histamine H1 receptor and meclizine or levocetirizine (a second-generation antihistamine) showed the conserved binding sites. This research illustrates the combined use of MicroED and molecular docking in unraveling protein-drug interactions for precision drug design and optimization.

---

Meclizine, marketed as “Antivert” or “Bonine”, is a first-generation H1 antihistamine used in the treatment of motion sickness and vertigo.^1-3^ Meclizine is chemically similar to other piperazine-class H1 antihistamines, such as cyclizine, buclizine, cetirizine, hydroxyzine, levocetirizine, and quetiapine.^4^ It consists of phenyl, chlorophenyl, and piperazine groups connected by a chiral carbon, with the methylbenzyl group linked on the other side of the piperazine ring (Figure 1A). The crystal structures of the piperazine-class antihistamines were mostly solved by single-crystal X-ray diffraction (SC-XRD) over the last several decades: cyclizine (1980),^5^ quetiapine (2005),^6^ cetirizine (2015),^7^ buclizine (2020).^8^ The disordered hydroxyzine model was derived by powder X-ray diffraction (PXRD) in 2019.^9^ The conventional SC-XRD encounters difficulties in obtaining large crystals from powdery substances,^10^ and solving PXRD structure can be challenging due to peak overlapping and broadening,^11^ therefore certain challenging crystal structures of piperazine-class H1 antihistamines were left unattainable for decades. The structure of meclizine remained elusive for more than 70 years despite its widespread medical use, ranking 142 as the most prescribed medicines in 2020 with more than 4 million prescriptions.^15^

**Figure 1.**
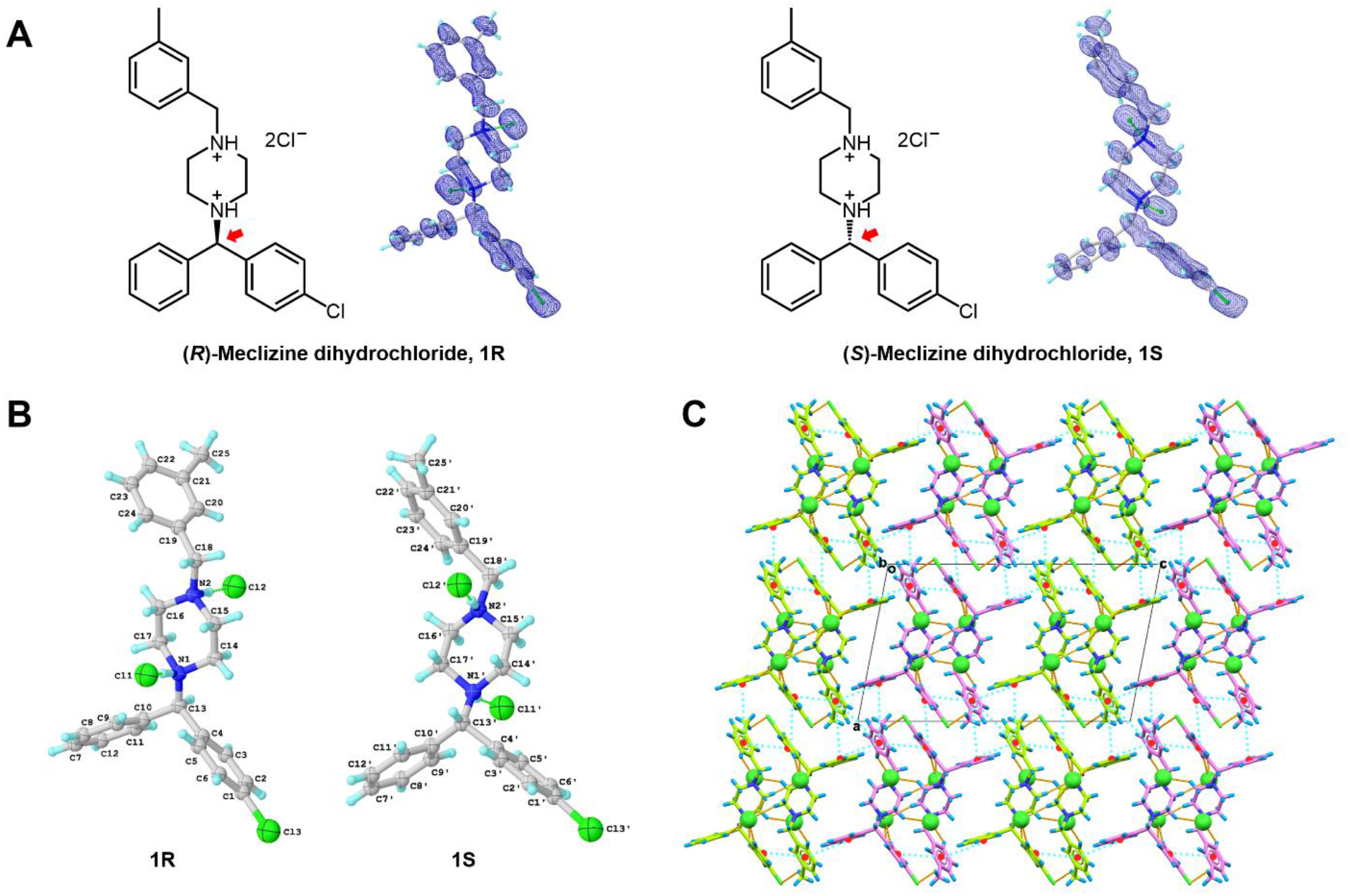
(A) Chemical structure and 2F_o_-F_c_ map (0.5 e·Å^-3^) of meclizine dihydrochloride (**1R**/**1S**). (B) Chemical notations of **1R** and **1S. 1R** was labeled with atom type and numbers, and **1S** was labeled with atom type and primed numbers. (C) Crystal packing diagram of **1R/1S**, viewed along *b* axis. **1R** was highlighted in green, **1S** was highlighted in violet. Hydrogen bonding interactions were represented by the dashed lines in orange, pi-stacking interactions were represented by the dashed line in cyan. Cl^-^ anions were highlighted in spacefill style. See details in Figure 2, Tables S3-S4 in the Supporting Information.

Microcrystal electron diffraction (MicroED) has emerged as a revolutionary technique that overcame the crystal size limitations of SC-XRD.^12,13^ It enables the analysis of nanocrystals directly from seemingly amorphous powder, that are merely a billionth the size needed for SC-XRD. The advent of MicroED has provided an alternative route for the structure elucidation of antihistamines with previously unknown crystal structures. For example, MicroED recently succeeded in solving the structure of levocetirizine, a compound whose crystal structure was unknown for 16 years after its initial medical use.^14^

The histamine H1 receptor is a member of the rhodopsin-like G protein-coupled receptor (GPCR) family that presents in various tissues, including smooth muscle, endothelial cells, and neurons in the central nervous system (CNS).^16-18^ Activation of this receptor by its biological agonist (histamine) regulates allergic responses, while the H1 antihistamine drugs reduce the receptor’s activity by binding and blocking the histamine interaction as an inverse agonist.^16-18^ First-generation antihistamines like meclizine involve several non-selective interactions with other receptors in the CNS, causing various adverse effects, such as drowsiness, dry mouth, and fatigue.^19^ While second-generation antihistamines like levocetirizine minimized adverse effects by reducing brain penetration and increasing binding selectivity.^20^ In this study, we used MicroED to determine the atomic structure of meclizine dihydrochloride. Molecular docking was then employed to analyze the binding between meclizine and the histamine H1 receptor, revealing its antihistamine mechanism and conformational changes between the drug formulation state and its biologically active state.

The meclizine dihydrochloride sample preparation for MicroED followed the previously described procedure (see details in the Supporting Information).^21^ The grid containing the crystals was examined using the 200 kV Thermo Fisher Talos Arctica Cryo-TEM with approximately 0.0251 Å wavelength. The microscope was equipped with a CetaD CMOS camera and EPUD software.^22^ The crystal thickness played a crucial role in obtaining optimal diffraction, so crystals were initially screened using imaging mode (LM 210×) for a grid atlas. Only crystals with a certain brightness contrast were thin enough and were selected for further analysis (Figure S1 in the Supporting Information). The eucentric height for each crystal was manually calibrated at low magnification (SA 3400×) to ensure proper centering during the continuous rotation. For data collection, a 70 µm C2 aperture and a 50 µm selected area (SA) aperture were utilized to reduce background noise and achieve a near parallel 1.4 µm beam size. The crystal was found to be very sensitive to radiation damage, *e*.*g*. crystal lattice was damaged after 40s even using the weakest spot size 11 under microprobe mode (0.0098 e^-1^/Å^2^/s), therefore an increased rotation rate of approximately 2 °/s over a smaller angular range of 80° (-40° to +40°), with an exposure time of 0.5 second per frame were used in order to minimize the total electron doses to 0.39 e^-1^/Å^2^. The MicroED movies were converted from mrc format to smv format using the mrc2smv software (available freely at https://cryoem.ucla.edu/microed).^22^ High-quality datasets were indexed and integrated using XDS,^23,24^ resulting in a completeness of over 55% for each dataset (Table S1 in the Supporting Information). The completeness was increased to 80.7% after scaling and merging data from two individual crystals (Table S2 in the Supporting Information). The intensities were converted to SHELX hkl format using XDSCONV.^24^ The MicroED structure was solved *ab initio* using SHELXT^25^ at a resolution of 0.96 Å. The MicroED structure of meclizine dihydrochloride contains two enantiomers, designated as **1R**/**1S**, was determined to be a centrosymmetric monoclinic space group P2_1_/c, with the unit cell of **a**=14.39 Å, **b**=7.19 Å, **c**=24.52 Å, ***α***=90.000°, ***β***=101.958°, ***γ***=90.000°. Subsequent refinement using SHELXL^26^ yielded a final R_1_ value of 17.89% (for more refinement statistics please see Table S2 in the Supporting Information). The positions of heavier atoms were accurately determined from the charge density map (Figure 1A). Since not all hydrogen (H) atoms could be located at this resolution, their positions were refined using a combination of constrained and free approaches.

The two enantiomers,**1R** and **1S** (See notations in Figure 1B), are packed as repetitive double layers (**1R**-**1R** or **1S**-**1S**) in the crystal lattice (Figure 1C). In the *b* axis, those layers are strengthened by various internal hydrogen bonding. Using the **1R**-**1R** layers for example, hydrogen bond interactions can be categorized into three groups (Figure 2A and Table S3 in the Supporting Information): (1) Two charge-assisted hydrogen bonds N1-H···Cl1^-^ and N2-H···Cl2^-^ are along the *b* axis, with distances at approximately 3.0 Å. (2) Seven C-H···Cl1^-^ and three C-H···Cl2^-^ hydrogen bonds that formed around Cl^-^ anions are along the *a* and *b* axes. These weak C-H···Cl^-^ hydrogen bonds, around 3.5-3.7 Å, were found between Cl^-^ anions and atoms in aromatic phenyl rings (C5, C9, C3, C24), piperazine rings (C14, C15), and alkane chains (C13, C18) of four surrounding **1R** molecules.^27^ (3) A weak C25-H···Cl3 hydrogen bond is established between methylbenzyl ring (C25) and chlorophenyl ring (Cl3).^27^ The same hydrogen bond geometry was found in **1S**-**1S** layers, albeit with different symmetries (Figure 2B and Table S3 in the Supporting Information). The packing along *a* and *c* axes is maintained by numerous pi-stacking interactions within the lattice, where three aromatic phenyl rings in a single **1R** or **1S** molecule can interact with up to fourteen aromatic rings from the surrounding ten molecules. The chlorophenyl ring 1 in **1R** for example interacts with rings 4-7 in parallel-displaced mode, the phenyl ring 2 interacts with rings 8-10 in T-shaped mode, and the methylbenzyl ring 3 interacts with rings 11-17 in a combination of parallel-displaced or T-shaped mode (Figures 2C and Table S4 in the Supporting Information).^28^ The identical pi-stacking geometry was observed in **1S** but interacting with different molecules (Figures 2D and Table S4 in the Supporting Information). Such interactions reinforce the packing within the **1R**-**1R** or **1S**-**1S** layers and establish connections between **1R** and **1S** molecules, further extending the crystal packing along the *a* and *c* axes. However, most of these interactions are remarkably weak, resulting in a fragile crystal lattice susceptible to external forces. ^27,28^

**Figure 2.**
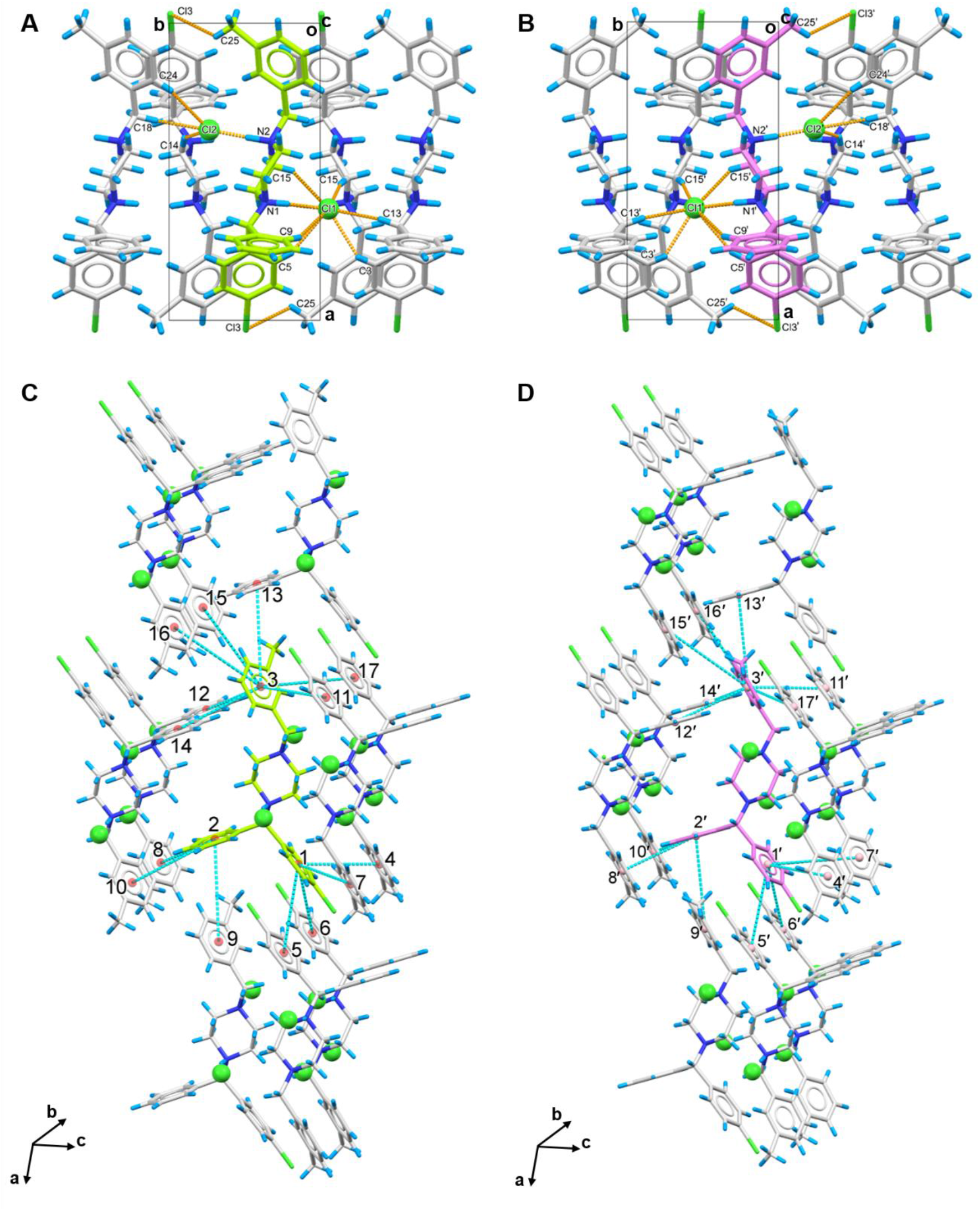
Hydrogen bonding and pi-stacking interactions in meclizine dihydrochloride (**1R/1S**) crystal packing. (A-B) Hydrogen bonding interactions in **1R** and **1S**, respectively, viewed along *c* axis; (C-D) Pi-stacking interactions in **1R** and **1S**, respectively, showing strong and moderate interactions with ten molecules in the surrounding. **1R** was colored in green, **1S** was colored in violet. Hydrogen bonding interactions were represented by the dashed lines in orange, pi-stacking interactions were represented by the dashed line in cyan. Cl^-^ anions were highlighted in spacefill style.

Like the related piperazine-class antihistamines, meclizine dihydrochloride (**1R**/**1S**) contains phenyl, chlorophenyl and piperazine rings which are connected by a chiral carbon, and the methylbenzyl ring is linked to the other side of piperazine ring (Figure 1). The crystal structure of **1R**/**1S** depicts the structure in its drug formulation state, serving as the initial reference prior for its transition into a biologically active conformation. Inspection of **1R**/**1S** showed similar C-C bond lengths without any substantial stretch or compression, varying from 1.47 Å to 1.57 Å (except phenyl rings), and the C-N bond lengths range from 1.40 Å to 1.55 Å. The C-C-N or C-N-C bond angles maintain a nearly perfect sp^3^ geometry, with an average value of 112.0 ± 4.0° in **1R**/**1S**. As for the heterocyclic piperazine rings, the N1-C14-C15-N2/N1′-C14′-C15′-N2′ and N1-C16-C17-N2/N1′-C16′-C17′-N2′ torsion angles are ± 57.8° and ± 63.7° in **1R**/**1S**, respectively. The distances between N1/N1′ and N2/N2′ atoms to the mean plane of C14-C15-C16-C17/C14′-C15′-C16′-C17′ are 0.72 Å and 0.65 Å, respectively. The piperazine rings maintain an almost perfect chair conformation due to the near 60° N-C-C-N torsion angles and comparable distances between the C-C-C-C planes and nitrogen atoms. The same conformation was also observed in other piperazine-class antihistamine structures such as buclizine monohydrochloride monohydrate (CSD entry: HUQVAT),^8^ and levocetirizine dihydrochloride (CSD entry: KIMDOD),^14^ suggesting the rigid conformation of piperazine ring which is unaffected by different charge states.

In **1R**/**1S**, five bonds corresponding to torsion angles *α* (N1-C13-C10-C9/ N1′-C13′-C10′-C9′), *β* (N1-C13-C4-C5/N1′-C13′-C4′-C5′), *θ* (C10-C13-N1-C17/C10′-C13′-N1′-C17′), *ω*(N1-C13-C10-C9/N1′-C13′-C10′-C9′) and *γ* (C13-C10-C9-C20/ C13′-C10′-C9′-C20′) have a relatively high degree of rotational freedom and can significantly influence the overall molecular conformation (Figure S2 in the Supporting Information). The orientation of phenyl and chlorophenyl rings is determined by the *α* and *β* torsion angles, which are approximately ± 47° (staggered-like), the piperazine ring was controlled by both *θ* and *ω* torsion angles, with values around ± 62° and ± 64° (staggered). The *γ* torsion angle is ± 79°, which manifested a staggered-like conformation and positioned the methylbenzyl ring to the direction of the phenyl group (Figure S2 in the Supporting Information). As previously mentioned, these torsions may be critical structural parameters in the drug formulation state but may not be important in its biologically active state upon interaction with the receptor. The latter may require a substantial conformational change.

The H1 antihistamine **1R**/**1S** acts as an inverse agonist, which inhibits the interaction between the agonist (histamine) and the histamine H1 receptor.^16-18^ However, the structural details of their binding mechanism remained unclear. Molecular docking of histamine H1 receptor in complex with **1R** or **1S** helped uncover the binding mechanism and the conformation changes between the drug formulation state and receptor-bound biologically active state of meclizine. A comparative study of the docking complexes between the receptor and **1R, 1S**, or levocetirizine (a second-generation antihistamine) identified a conserved binding sites. In the setup for molecular docking, the enantiomerically pure ligand structure of **1R, 1S**, or levocetirizine (CSD entry: KIMDOD) ^14^ was directly obtained from their MicroED structures, with the polar H atoms removed (see details in the Supporting Information). All active torsion angles, including *α, β, θ, ω*, and *γ*, were rendered rotatable during the docking. The cryo-EM structure of the histamine H1 receptor (PDB entry: 7DFL) was used as a rigid model.^29^ The molecular docking was conducted by AutoDock Vina 1.1.2^30,31^ using an 18.75 Å × 18.75 Å × 18.75 Å grid box with 0.375 Å spacing, centered at the experimentally determined ligand (histamine) position (see Figure S3 in the Supporting Information).

The cryo-EM structure of the complex formed between the histamine H1 receptor and its agonist (histamine) revealed interactions within four specific transmembrane helices (TMs): **I, II, III** and **V**. These interactions involved: (1) Two weak hydrogen bonds between Asp107 and Tyr458 residues and histamine’s ethylamine group; (2) Three strong hydrogen bonds between Thr112, Asn198, Tyr431 and histamine’s imidazole ring (see Figure 3A, Table S5 in the Supporting Information).

**Figure 3.**
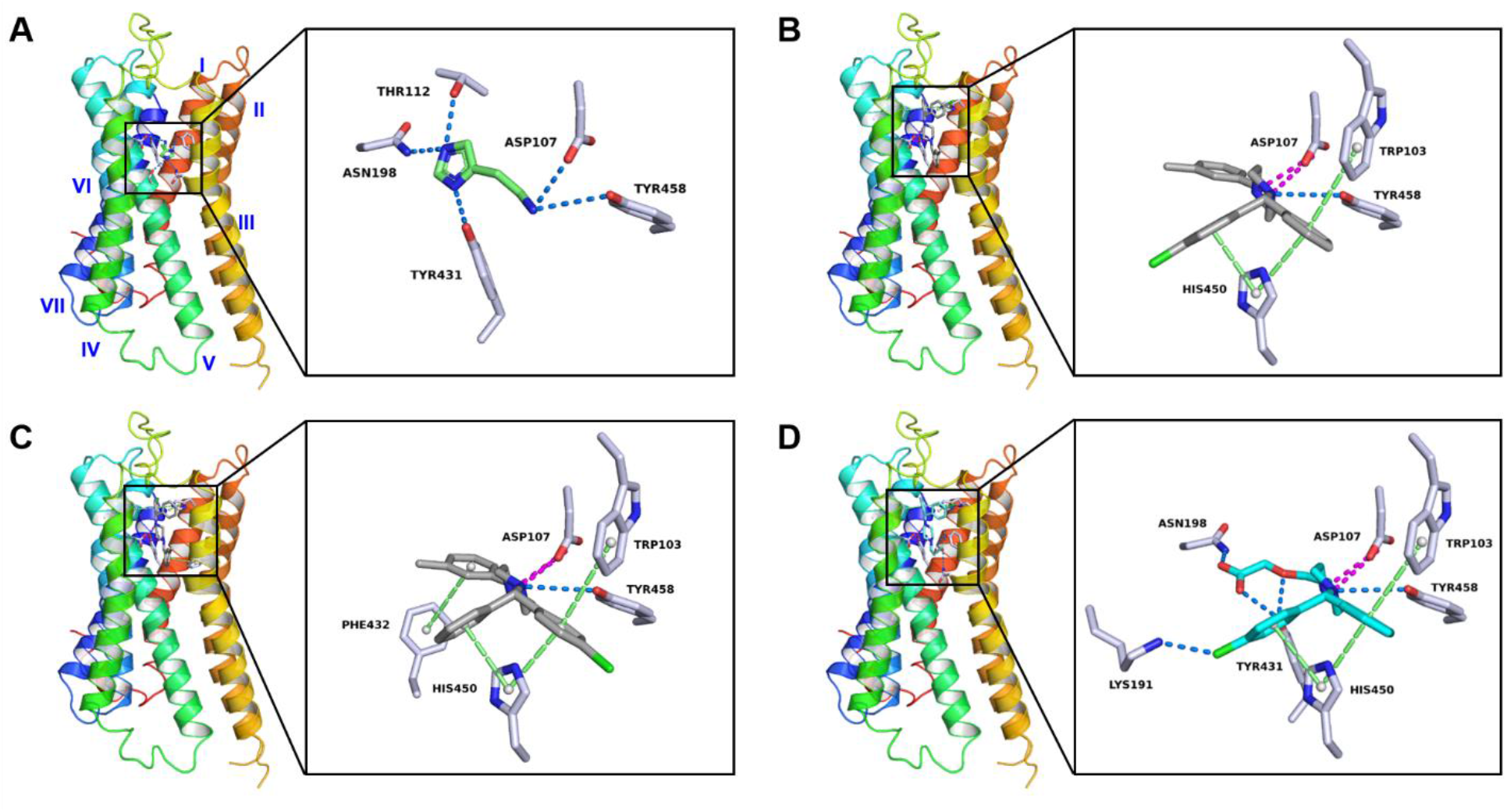
Protein–drug interaction diagram of complex between histamine H1 receptor and (A) histamine, (B) **1R**, (C) **1S** and (D) levocetirizine. Histamine was colored in green, **1R** and **1S** were colored in grey, and levocetirizine was colored in light blue. Hydrogen bonding and halogen bonding interactions were colored by the dashed line in marine, pi-stacking and pi-cation interactions were colored by the dashed line in lime, and salt bridges were colored by the dashed line in magenta. Seven transmembrane *α*-helical segments (**I**-**VII**) in histamine H1 receptor were colored from red to blue, see the overall view in Figure S3 in the Supporting Information.

The molecular docking analysis of the histamine H1 receptor complexed with **1R** or **1S** involved some of the same histamine binding residues but also suggested additional binding residues at adjacent regions. The major interactions between the receptor and **1R** involve residues from transmembrane helices (TMs) **I, V** and **VI**, including (1) One or two salt bridges between Asp107 and the protonated piperazine ring in **1R**; (2) Weak hydrogen bond between Tyr458 and the piperazine ring in **1R**; (3) Pi-stacking interactions involving His450, Trp103, and the phenyl or chlorophenyl ring in **1R**; (4) Up to seven hydrophobic interactions involving Tyr, Ile, Trp, and Phe residues in TMs **III** and **V**, stabilizing the orientation of the methylbenzyl ring towards the left. These conformation changes are mainly driven by the rotation of the *θ* torsion angle by more than 100°, resulting in the piperazine ring being nearly perpendicular to the plane formed by phenyl and chlorophenyl rings (Figure S2 in the Supporting Information). The enantiomer **1S** displayed interactions similar to **1R**, with an additional pi-stacking interaction between Phe432 and the methylbenzyl ring, possibly enhancing its binding affinity. These conformational shifts were led primarily by rotations of the θ, ω, γ torsions in **1S**, reorienting the phenyl and chlorophenyl rings, yet maintaining the piperazine and methylbenzyl rings in a position analogous to **1R** (Figure S2 in the Supporting Information).

Levocetirizine, a second-generation antihistamine, exhibits enhanced binding selectivity and minimal brain penetration compared to **1R**/**1S**.^32^ The molecular docking analysis of the histamine H1 receptor complexed with levocetirizine showed the conserved binding sites of TRP103, Asp107, His450, and Tyr458 which were consistent with those for **1R**/**1S** (Figure 3D). However, the binding interactions with levocetirizine were stronger and more closely resembled histamine than **1R**/**1S**. For example, the strong hydrogen bonds between Asn198, Tyr431 and the levocetirizine’s ethoxyacetic acid group; The robust halogen bond between Lys191 and the levocetirizine’s chlorophenyl group, which has been validated in the biochemical binding analysis in the literature.^33^

In summary, the 3D structure of meclizine dihydrochloride (**1R**/**1S**) was determined directly from seemingly amorphous powders by MicroED. The R and S enantiomers packed as repetitive double layers (**1R**-**1R** or **1S**-**1S**) in the lattice, facilitated by robust N-H···Cl^-^ hydrogen bonding interactions and weaker interactions like C-H···Cl^-^ hydrogen bonds and pi-stacking. Molecular docking revealed the binding mechanism of meclizine to the histamine H1 receptor identifying differences between its drug formulation structure and its biologically active receptor bound structure. A comparison of the docking complexes between histamine H1 receptor and meclizine or levocetirizine revealed conserved binding sites at Trp103, Asp107, His450 and Tyr458 within the receptor. The new structural information established a basis for understanding binding mechanism of H1 antihistamines and receptor, using a combined approach of MicroED and molecular docking which would be useful for future precision drug design and optimization pipelines.

## Acknowledgements

The authors thank Michael W. Martynowycz for support and discussions. This study was funded in part by the National Institutes of Health P41GM136508. Portions of this research or manuscript completion were developed with funding from the Department of Defense grants MCDC-2202-002. Effort sponsored by the U.S. Government under Other Transaction number W15QKN-16-9-1002 between the MCDC, and the Government. The US Government is authorized to reproduce and distribute reprints for Governmental purposes, notwithstanding any copyright notation thereon. The views and conclusions contained herein are those of the authors and should not be interpreted as necessarily representing the official policies or endorsements, either expressed or implied, of the U.S. Government. The PAH shall flowdown these requirements to its subawardees, at all tiers. The Gonen laboratory is supported by funds from the Howard Hughes Medical Institute.

