## Supplemental Information for "Unraveling the Structure of Meclizine Dihydrochloride with MicroED"

### Methods

#### Materials.

Meclizine dihydrochloride, (*R/S*)-1-[(4-chlorophenyl)(phenyl)methyl]-4-(3-methylbenzyl)piperazine, was commercially purchased from InvivoChem and used as received without further recrystallization.

#### Grid preparation.

Sample preparation followed the procedure as described previously.[1] One carbon-coated copper grid (400-mesh, 3.05 mm O.D., Ted Pella Inc.) was pretreated with glow-discharge plasma at 15 mA on the negative mode using PELCO easiGlow (Ted Pella Inc.) for 60s. Around 1 mg of powdery compounds were carefully weighed by a Mettler Toledo (XPR225DR) analytical balance and mixed with a grid in a 10 mL scintillation vial. After gently shaking the vial, the grid was removed and clipped using c-ring and autogrid clip (Thermo Fisher) at room temperature.

#### MicroED data collection.

The clipped grid was loaded in an aligned Thermo Fisher Talos Arctica Cryo-TEM (200 kV, ~0.0251 Å) at 100 K, equipped with a CetaD CMOS camera (4096 × 4096 pixels) and EPUD (Thermo Fisher) software.[1,2] Screening of size- and thickness-suitable microcrystals was done in the imaging mode (LM 210× and SA 3400×). The MicroED data was collected in the diffraction mode with 659 mm diffraction length (the calibrated sample-detector distance), 70 µm C2 aperture, and a 50 µm selected area (SA) aperture in the parallel beam condition (45.2% C2 intensity) which resulted in a beam size at approximately 1.4 µm. Typical data collection used a constant rotation rate of ~2° per second over an angular wedge of 80° from -40° to +40°, with 0.5s exposure time per frame. Crystal selected for MicroED data collection were isolated and calibrated to eucentric heights to maintain the crystals inside the beam during the rotation.

**MicroED data processing.**

The MicroED data was saved in mrc format and converted to smv format using the mrc2smv software (https://cryoem.ucla.edu/microed).[2] The converted frames were indexed and integrated by XDS.[3,4] Two selected datasets with the highest resolution were scaled and merged using XSCALE,[4] and intensities were converted to SHELX hkl format using XDSCONV.[4] The merged dataset showed 80.7% overall completeness, which can be *ab initio* solved by SHELXT[5] at a resolution of 0.96 Å (Table S1). The structure was refined by SHELXL[6] in Shelxle[7] as a graphical interference to yield the final MicroED structure (Figure 1 and Table S2).

**Molecular Docking.**

The ligand structures of **1R/1S** were extracted from the refined MicroED structure and saved as mol2 files. The structure of levocetirizine was downloaded from CSD database (CSD entries: KIMDOD) and transformed to mol2 files. The $Cl^-$ anions, water molecules, and polar hydrogen atoms from amine in the piperazine ring (pKa ≈ 2.12 and 6.55)[8] or ethoxyacetate group (pKa ≈ 2.9)[9] were removed. Then the ligand structures were imported into AutoDock Tools 1.5.7,[10] where all active torsion bonds were made rotatable.

The Cryo-EM structure of histamine H1 receptor (PDB entry: 7DFL)[11] was retrieved from the Protein Data Bank (https://www.rcsb.org/) after removing the ligand and other protein molecules using Pymol 2.5.5.[12] Subsequently, hydrogen atoms and charges were computed and added using AutoDock Tools 1.5.7.[10]

A grid box measuring 18.75 Å × 18.75 Å × 18.75 Å with 0.375 Å spacing was positioned along the *x*-, *y*-, and *z*-axes. The grid center was determined based on the experimental ligand (histamine) position at coordinates (131.312, 132.360, 158.503), see Figure S3.

The AutoDock Vina 1.1.2 was used for molecular docking,[13,14] where all active torsion bonds in ligands were set to be flexible, and the receptor was set as a rigid model. The docked complex with the minimum binding energy was exported and analyzed by the Protein-Ligand Interaction Profiler (PLIP) web tool and Pymol 2.5.5 (Figure 3 and Tables S5-S8).[12,15]

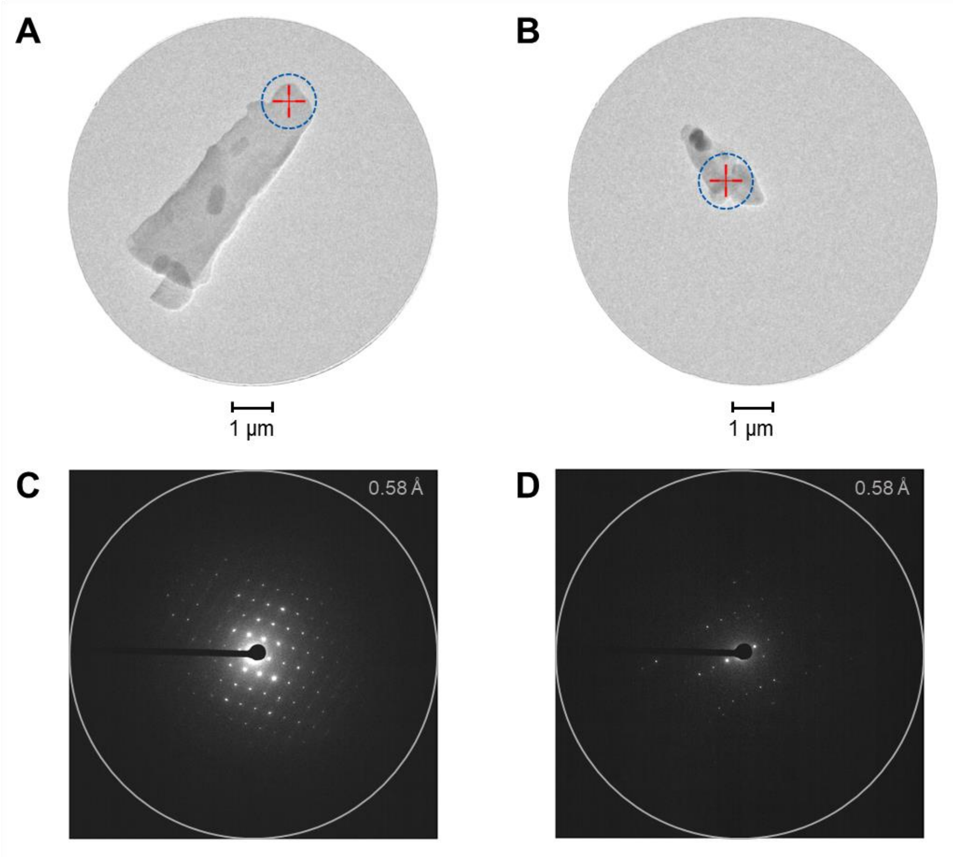


**Figure S1** Crystal appearance and diffraction pattern under the TEM. (A-B) Images of items 1 and 2 under the imaging mode (SA 3400×), respectively. The diffraction beam size was highlighted in dashed blue circles; (C-D) Diffraction pattern of items 1 and 2 under diffraction mode (659 mm), respectively. The integration edge was colored in grey rings.

**A**

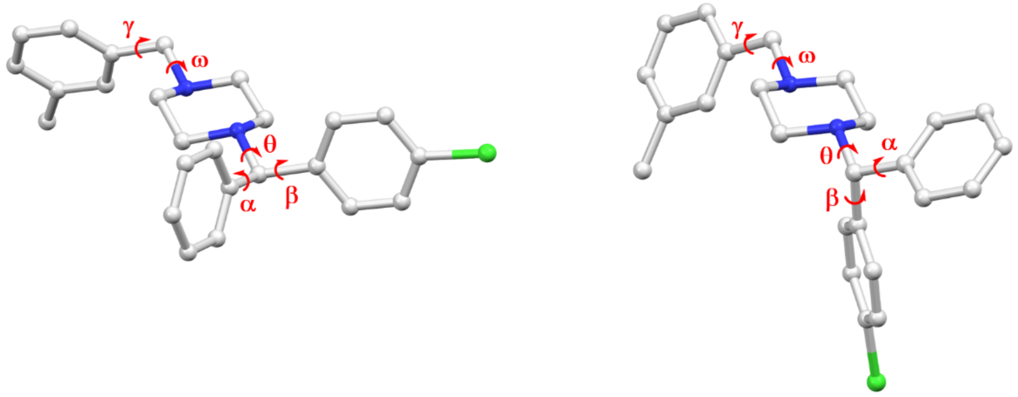


| **Crystal Structure:** | |
|---|---|
| α (N1-C13-C10-C9) | -46.44$^{o}$ |
| β (N1-C13-C4-C5) | +47.58$^{o}$ |
| θ (C10-C13-N1-C17) | -61.68$^{o}$ |
| ω (C16-N2-C18-C19) | -64.46$^{o}$ |
| γ (N2-C18-C19-C20) | -79.05$^{o}$ |

| **Molecular Docking:** | |
|---|---|
| α (N1-C13-C10-C9) | -13.55$^{o}$ |
| β (N1-C13-C4-C5) | +2.42$^{o}$ |
| θ (C10-C13-N1-C17) | -170.24$^{o}$ |
| ω (C16-N2-C18-C19) | -83.66$^{o}$ |
| γ (N2-C18-C19-C20) | -6.60$^{o}$ |

**B**

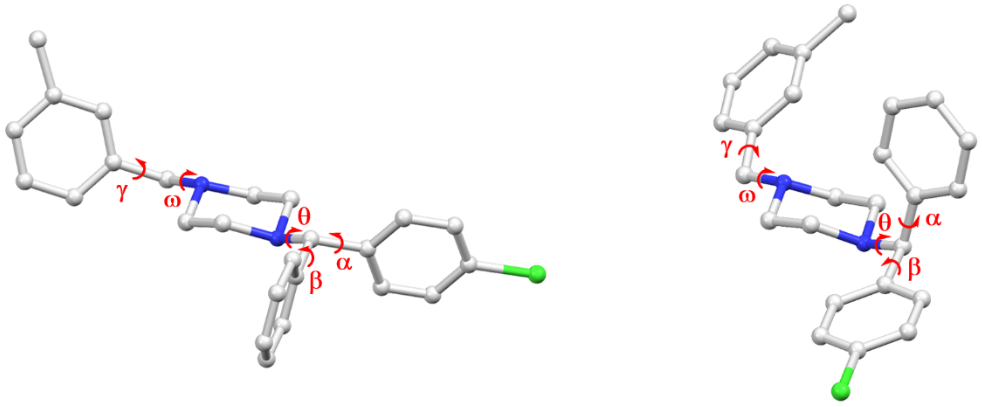


| **Crystal Structure:** | |
|---|---|
| α (N1'-C13'-C10'-C9') | +46.44$^{o}$ |
| β (N1'-C13'-C4'-C5') | -47.58$^{o}$ |
| θ (C10'-C13'-N1'-C17') | +61.68$^{o}$ |
| ω (C16'-N2'-C18'-C19') | +64.46$^{o}$ |
| γ (N2'-C18'-C19'-C20') | +79.05$^{o}$ |

| **Molecular Docking:** | |
|---|---|
| α (N1'-C13'-C10'-C9') | +15.36$^{o}$ |
| β (N1'-C13'-C4'-C5') | -13.75$^{o}$ |
| θ (C10'-C13'-N1'-C17') | -65.02$^{o}$ |
| ω (C16'-N2'-C18'-C19') | +169.99$^{o}$ |
| γ (N2'-C18'-C19'-C20') | -4.76$^{o}$ |

**Figure S2** The major conformation changes between the crystal structures of **1R/1S** and their molecular docking structures. See notations in Figure 1B.

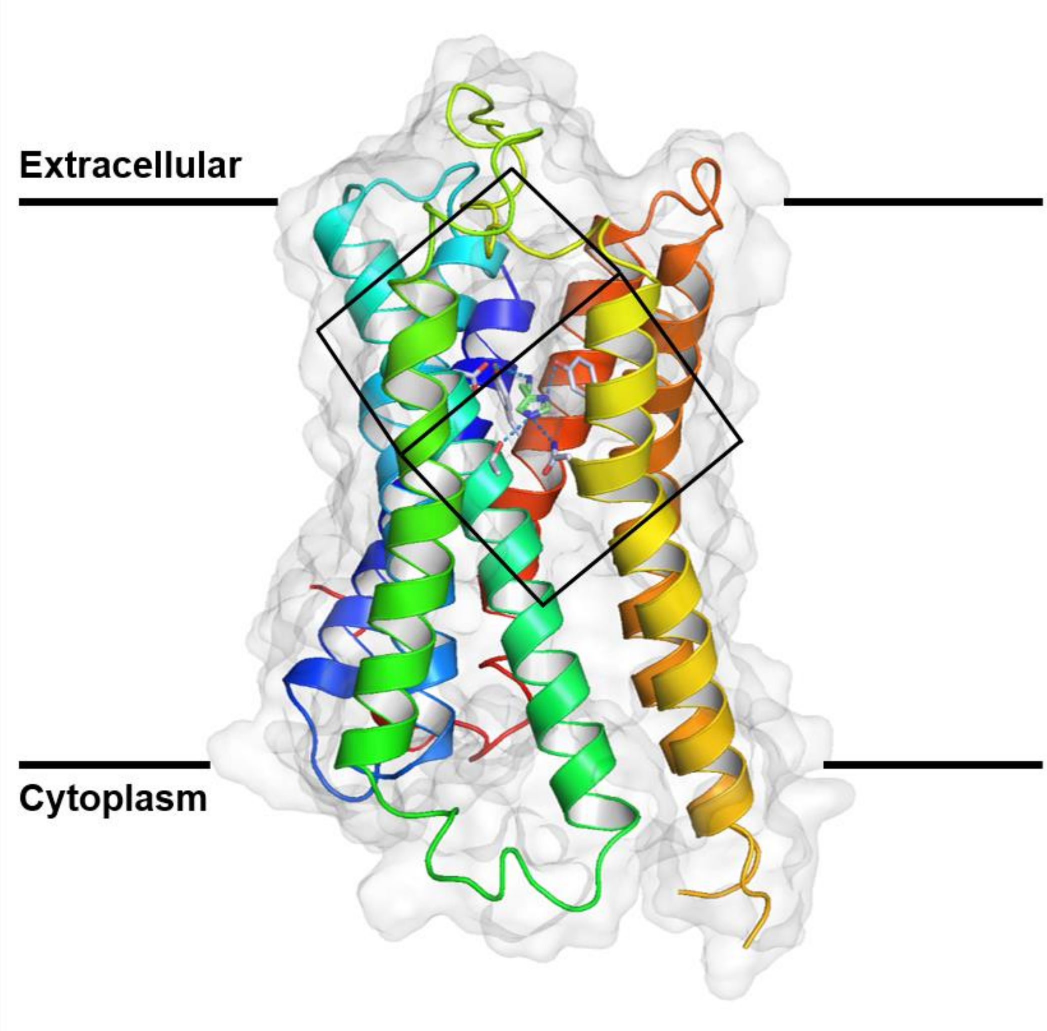


**Figure S3** Overall view of the complex between histamine H1 receptor and histamine determined by Cryo-EM (PDB entry: 7DFL). The 18.75 Å × 18.75 Å × 18.75 Å grid box used for molecular docking was centered by the experimental ligand (histamine) position at coordinates (131.312, 132.360, 158.503), marked by black lines.

**Table S1** MicroED data statistics of two selected items of meclizine dihydrochloride.

| | Item 1 | Item 2 |
|---|---|---|
| Space group | $P2_1$ | $P2_1$ |
| Unit cell lengths (Å) | | |
| a | 14.39 | 14.00 |
| b | 7.19 | 7.17 |
| c | 24.52 | 24.41 |
| Unit cell angles (°) | | |
| $\alpha$ | 90.000 | 90.000 |
| $\beta$ | 100.958 | 103.276 |
| $\gamma$ | 90.000 | 90.000 |
| No. of observed reflections | 4870 | 4845 |
| No. of unique reflections | 1869 | 1812 |
| $R_{obs}$ (%) | 12.7 | 22.6 |
| $R_{meas}$ (%) | 16.1 | 28.6 |
| I/Sigma | 5.14 | 3.51 |
| $CC_{1/2}$ | 99.3 | 96.9 |
| Resolution (Å) | 0.95 | 0.95 |
| Completeness (%) | 55.8 | 56.3 |

**Table S2** MicroED data statistics of meclizine dihydrochloride (merged).

| | |
|---|---|
| Stoichiometric formula | $C_{25}H_{29}Cl_3N_2$ |
| Mr | 463.85 |
| Temperature (K) | 100 |
| Crystal system | Monoclinic |
| Space group | $P2_1/c$ |
| Unit cell lengths (Å) | |
| a | 14.39 |
| b | 7.19 |
| c | 24.52 |
| Unit cell angles (°) | |
| $\alpha$ | 90.000 |
| $\beta$ | 101.958 |
| $\gamma$ | 90.000 |
| Cell volume ($Å^3$) | 2493.49 |
| No. of observed reflections | 9487 |
| No. of unique reflections | 2623 |
| $R_{obs}$ (%) | 19.0 |
| $R_{meas}$ (%) | 22.3 |
| I/Sigma | 3.81 |
| $CC_{1/2}$ | 98.9 |
| Resolution (Å) | 0.96 |
| Completeness (%) | **80.7** |
| $R_1$ (%) | **17.89** |
| $wR_2$ (%) | 42.43 |
| GooF | 1.737 |

**Table S3** Hydrogen-bond geometry in meclizine dihydrochloride **1R/1S** (Å, °).

| **R form** | **D–H** | **H···A** | **D···A** | **D–H···A** | **Type** |
|---|---|---|---|---|---|
| N1–H···Cl1 | 1.008 | 2.041 | 3.013 | 161.27 | N–H···Cl$^-$ |
| N2–H···Cl2 | 1.117 | 1.877 | 2.994 | 178.15 | |
| C3–H···Cl1$^{i}$ | 0.930 | 2.711 | 3.473 | 139.75 | C–H···Cl$^-$ |
| C5–H···Cl1 | 0.930 | 2.804 | 3.679 | 157.28 | |
| C9–H···Cl1 | 0.930 | 2.726 | 3.593 | 155.60 | |
| C13–H···Cl1$^{i}$ | 0.981 | 2.677 | 3.598 | 156.45 | |
| C15–Ha···Cl1$^{ii}$ | 0.969 | 2.754 | 3.709 | 168.59 | |
| C15–Hb···Cl1 | 0.971 | 2.933 | 3.647 | 131.29 | |
| C14–H···Cl2$^{iv}$ | 0.969 | 2.765 | 3.569 | 140.75 | |
| C18–H···Cl2$^{iii}$ | 0.972 | 2.656 | 3.605 | 165.30 | |
| C24–H···Cl2$^{iii}$ | 0.931 | 2.900 | 3.719 | 147.42 | |
| C25–H···Cl3$^{ii}$ | 0.961 | 2.921 | 3.570 | 125.90 | C–H···Cl |
| **S form** | **D–H** | **H···A** | **D···A** | **D–H···A** | **Type** |
| N1′–H···Cl1′ | 1.008 | 2.041 | 3.013 | 161.27 | N–H···Cl$^-$ |
| N2′–H···Cl2′ | 1.117 | 1.877 | 2.994 | 178.15 | |
| C3′–H···Cl1′$^{iii}$ | 0.930 | 2.711 | 3.473 | 139.75 | C–H···Cl$^-$ |
| C5′–H···Cl1′ | 0.930 | 2.804 | 3.679 | 157.28 | |
| C9′–H···Cl1′ | 0.930 | 2.726 | 3.593 | 155.60 | |
| C13′–H···Cl1′$^{iii}$ | 0.981 | 2.677 | 3.598 | 156.45 | |
| C15′–Ha···Cl1′$^{v}$ | 0.969 | 2.754 | 3.709 | 168.59 | |
| C15′–Hb···Cl1′ | 0.971 | 2.933 | 3.647 | 131.29 | |
| C14′–H···Cl2′$^{vi}$ | 0.969 | 2.765 | 3.569 | 140.75 | |
| C18′–H···Cl2′$^{i}$ | 0.972 | 2.656 | 3.605 | 165.30 | |
| C24′–H···Cl2′$^{i}$ | 0.931 | 2.900 | 3.719 | 147.42 | |
| C25′–H···Cl3′$^{ii}$ | 0.961 | 2.921 | 3.570 | 125.90 | C–H···Cl |
| Symmetry codes: (i) x,-1+y,z; (ii) 1-x,-1/2+y,1.5-z; (iii) x,1+y,z; (iv) 1-x,1/2+y,1.5-z; (v) 1-x,1/2+y,1/2-z; (vi) 1-x,-1/2+y,1/2-z. | | | | | |

**Notes**: See details in Figure 2.

**Table S4** Pi-stacking interactions in meclizine dihydrochloride **1R/1S** (Å, °).

| ***pi-stacking interactions in 1R*** | | | | |
|---|---|---|---|---|
| **Centroid 1**[a] | **Centroid 2**[a] | **Distance**[b] | **Relative Orientation**[c] | **Type** |
| 1 | 4 | **4.58** | 19.22 | Parallel-displaced |
| | 5 | 5.65 | 12.57 | Parallel-displaced |
| | 6 | 5.65 | 12.57 | Parallel-displaced |
| | 7 | 6.6 | 19.22 | Parallel-displaced |
| 2 | 8 | **4.85** | 74.69 | T-shaped |
| | 9 | 5.59 | 74.69 | T-shaped |
| | 10 | 6.09 | 74.69 | T-shaped |
| 3 | 11 | **4.58** | 19.22 | Parallel-displaced |
| | 12 | **4.85** | 74.69 | T-shaped |
| | 13 | 5.59 | 74.69 | T-shaped |
| | 14 | 6.09 | 74.69 | T-shaped |
| | 15 | 6.50 | 0 | Parallel-displaced |
| | 16 | 6.59 | 0 | Parallel-displaced |
| | 17 | 6.60 | 19.22 | Parallel-displaced |
| ***pi-stacking interactions in 1S*** | | | | |
| **Centroid 1**[a] | **Centroid 2**[a] | **Distance**[b] | **Relative Orientation**[c] | **Type** |
| 1′ | 4′ | **4.58** | 19.22 | Parallel-displaced |
| | 5′ | 5.65 | 12.57 | Parallel-displaced |
| | 6′ | 5.65 | 12.57 | Parallel-displaced |
| | 7′ | 6.6 | 19.22 | Parallel-displaced |
| 2′ | 8′ | **4.85** | 74.69 | T-shaped |
| | 9′ | 5.59 | 74.69 | T-shaped |
| | 10′ | 6.09 | 74.69 | T-shaped |
| 3′ | 11′ | **4.58** | 19.22 | Parallel-displaced |
| | 12′ | **4.85** | 74.69 | T-shaped |
| | 13′ | 5.59 | 74.69 | T-shaped |
| | 14′ | 6.09 | 74.69 | T-shaped |
| | 15′ | 6.50 | 0 | Parallel-displaced |
| | 16′ | 6.59 | 0 | Parallel-displaced |
| | 17′ | 6.60 | 19.22 | Parallel-displaced |

**Notes**: [a]Centroids were determined as the center of aromatic phenyl rings in **1R**/**1S**. [b]Distances were measured between centroids.[c]Relative orientations were measured by the angles between planes of two aromatic rings. See details in Figure 2.

**Table S5** Protein-ligand interactions of histamine H1 receptor and histamine determined by Cryo-EM (Å, °).

| **Hydrogen Bonds** | | | | | |
|---|---|---|---|---|---|
| Residue | AA | Ligand Group[a] | H–A | D–A | D-H···A |
| 107R | ASP | $EtNH_2$ | 2.41 | 3.38 | 158.95 |
| 458R | TYR | $EtNH_2$ | 3.16 | 3.94 | 139.55 |
| 112R | THR | Im | 2.35 | 3.07 | 127.05 |
| 198R | ASN | Im | 2.47 | 3.09 | 118.28 |
| 431R | TYR | Im | 2.18 | 3.02 | 147.81 |
| **Hydrophobic Interactions** | | | | | |
| Residue | AA | Ligand Group[a] | Distance | | |
| 108R | TYR | $EtNH_2$ | 3.75 | | |
| 431R | TYR | $EtNH_2$ | 3.82 | | |

**Notes**: [a]Ligand group was represented by abbreviation: imidazole (Im), ethylamine ($EtNH_2$).

**Table S6** Protein-ligand interactions of histamine H1 receptor and **1R** complex predicted by molecular docking (Å, °).

| | | | | | |
|---|---|---|---|---|---|
| **Salt Bridges** | | | | | |
| Residue | AA | Distance | Protein positive? | Ligand Group[a] | |
| 107R | ASP | 3.76 | No | Pip | |
| 107R | ASP | 3.69 | No | Pip | |
| **Hydrogen Bonds** | | | | | |
| Residue | AA | Ligand Group[a] | H–A | D–A | D–H···A |
| 458R | TYR | Pip | 3.30 | 4.00 | 132.70 |
| **π-Stacking** | | | | | |
| Residue | AA | Ligand Group[a] | Distance[b] | Angle[c] | Stacking Type |
| 103R | TRP | Ph | 4.62 | 66.31 | T-shaped |
| 450R | HIS | Ph | 4.03 | 71.31 | T-shaped |
| 450R | HIS | ClPh | 3.79 | 65.97 | T-shaped |
| **Hydrophobic Interactions** | | | | | |
| Residue | AA | Ligand Group[a] | Distance | | |
| 454R | ILE | Ph | 3.90 | | |
| 87R | TYR | Ph | 3.19 | | |
| 454R | ILE | Ph | 3.44 | | |
| 108R | TYR | ClPh | 3.94 | | |
| 182R | THR | ClPh | 2.56 | | |
| 108R | TYR | MeBn | 3.69 | | |
| 115R | ILE | MeBn | 3.62 | | |
| 158R | TRP | MeBn | 3.46 | | |
| 428R | TRP | MeBn | 3.33 | | |
| 428R | TRP | MeBn | 3.97 | | |
| 431R | TYR | MeBn | 3.47 | | |
| 432R | PHE | MeBn | 3.18 | | |

**Notes**: [a]Ligand group was represented by abbreviation: piperazine (Pip), phenyl (Ph), chlorophenyl (ClPh), methylbenzyl (MeBn). [b]Distances were measured between centroids determined as the center of aromatic rings. [c]Angles were measured by the angles between planes of two aromatic rings.

**Table S7** Protein-ligand interactions of histamine H1 receptor and **1S** complex predicted by molecular docking (Å, °).

| **Salt Bridges** | | | | | |
|---|---|---|---|---|---|
| Residue | AA | Distance | Protein positive? | Ligand Group[a] | |
| 107R | ASP | 3.68 | No | Pip | |
| 107R | ASP | 3.79 | No | Pip | |
| **Hydrogen Bonds** | | | | | |
| Residue | AA | Ligand Group[a] | H–A | D–A | D–H···A |
| 458R | TYR | Pip | 3.33 | 4.08 | 137.47 |
| **π-Stacking** | | | | | |
| Residue | AA | Ligand Group[a] | Distance[b] | Angle[c] | Stacking Type |
| 450R | HIS | Ph | 3.80 | 70.65 | T-shaped |
| 103R | TRP | ClPh | 4.74 | 65.54 | T-shaped |
| 450R | HIS | ClPh | 3.90 | 73.94 | T-shaped |
| 432R | PHE | MeBn | 4.83 | 70.17 | T-shaped |
| **Hydrophobic Interactions** | | | | | |
| Residue | AA | Ligand Group[a] | Distance | | |
| 108R | TYR | Ph | 3.84 | | |
| 182R | THR | Ph | 2.56 | | |
| 431R | TYR | Ph | 3.80 | | |
| 87R | TYR | ClPh | 3.45 | | |
| 454R | ILE | ClPh | 3.18 | | |
| 454R | ILE | ClPh | 3.69 | | |
| 108R | TYR | MeBn | 3.71 | | |
| 115R | ILE | MeBn | 3.69 | | |
| 158R | TRP | MeBn | 3.54 | | |
| 428R | TRP | MeBn | 3.84 | | |
| 428R | TRP | MeBn | 3.39 | | |
| 431R | TYR | MeBn | 3.25 | | |

**Notes**: [a]Ligand group was represented by abbreviation: piperazine (Pip), phenyl (Ph), chlorophenyl (ClPh), methylbenzyl (MeBn). [b]Distances were measured between centroids determined as the center of aromatic rings. [c]Angles were measured by the angles between planes of two aromatic rings.

**Table S8** Protein-ligand interactions of histamine H1 receptor and levocetirizine complex predicted by molecular docking (Å, °).

| | | | | | |
|---|---|---|---|---|---|
| **Salt Bridges** | | | | | |
| Residue | AA | Distance | Protein positive? | Ligand Group[a] | |
| 107R | ASP | 3.76 | No | Pip | |
| 107R | ASP | 3.73 | No | Pip | |
| **Hydrogen Bonds** | | | | | |
| Residue | AA | Ligand Group[a] | H–A | D–A | D–H···A |
| 198R | ASN | EtOAc | 2.25 | 2.98 | 126.87 |
| 431R | TYR | EtOAc | 2.28 | 3.10 | 144.14 |
| 431R | TYR | EtOAc | 1.92 | 2.82 | 158.91 |
| 458R | TYR | Pip | 3.28 | 4.03 | 136.96 |
| **π-Stacking** | | | | | |
| Residue | AA | Ligand Group[a] | Distance[b] | Angle[c] | Stacking Type |
| 103R | TRP | Ph | 4.54 | 71.57 | T-shaped |
| 450R | HIS | Ph | 4.08 | 71.58 | T-shaped |
| 450R | HIS | ClPh | 3.87 | 72.53 | T-shaped |
| **Halogen Bonds** | | | | | |
| Residue | AA | Ligand Group[a] | **X**···Y | A–**X**···Y | **X**···Y–B |
| 191R | LYS | ClPh | 3.02 | 140.96 | 138.51 |
| **Hydrophobic Interactions** | | | | | |
| Residue | AA | Ligand Group[a] | Distance | | |
| 87R | TYR | Ph | 3.22 | | |
| 454R | ILE | Ph | 3.78 | | |
| 454R | ILE | Ph | 3.95 | | |
| 108R | TYR | ClPh | 3.87 | | |
| 182R | THR | ClPh | 2.72 | | |
| 431R | TYR | ClPh | 3.61 | | |

**Notes**: [a]Ligand group was represented by abbreviation: piperazine (Pip), phenyl (Ph), chlorophenyl (ClPh), ethoxyacetic acid (EtOAc). [b]Distances were measured between aromatic ring centroids or centroid-cation. [c]Angles were measured by the angles between planes of two aromatic rings.
